# Is It There? - Estimating The Invasion of Armoured Sailfin Catfish (*Pterygoplichthys Sp*.) in the Water Bodies of Eastern Ghats, India Using eDNA Approach

**DOI:** 10.1101/2023.08.26.554971

**Authors:** Neeldeep Ganguly, Govindhaswamy Umapathy

## Abstract

Early detection of invasive species is crucial for effective control of the potential damage they can inflict on the ecosystems. In contrast to the many limitations that traditional detection methods like visual surveys and netting hold, the use of environmental DNA assay provides a powerful alternative. This non-invasive, highly sensitive, and user-friendly technique offers the advantage of detecting invasive species even in areas where direct observation is challenging, thus addressing the shortcomings of traditional techniques and enhancing overall accuracy in estimating distribution. The spread of invasive *Pterygoplichthys sp*. has become a cause for concern in biodiversity-rich countries like India. Despite this, comprehensive studies on the prevalence of this invasive species are limited. The Eastern Ghats of India remain underexplored with a high potential for supporting diverse lifeforms. Studying the extent of biological invasions in the Eastern Ghats is very essential for effective conservation management to mitigate the ecological and socioeconomic impacts of invasive species. In this study, we have designed and optimised an eDNA-based quantitative PCR assay to map the presence and spread of invasive *Pterygoplichthys sp*. in selected freshwater ecosystems of the Eastern Ghats. With this assay, we detected invasive *Pterygoplichthys sp* in almost 65% of the total locations sampled. This study can be further extended to larger geographical areas, which in turn can contribute in formulating necessary measures by the authorities to manage invasion and conserve the diversity of the freshwater ecosystem.

## 2. INTRODUCTION

It is astonishing how freshwater makes up only 2.5% of the Earth’s water, and only a tiny fraction of that is available as habitable freshwater. Despite this, freshwater ecosystems like rivers, lakes, and wetlands are home to over 40% of the world’s fish species (Hood, 2020). Freshwater ecosystems are more complex in terms of physical and chemical parameters than marine ecosystems, providing an enormous range of habitats and niches for fishes to occupy and thrive in. (Stanford, et al.,2005) This results in a high level of freshwater fish diversity compared to the amount of freshwater available to us, which is often referred to as the “Freshwater fish paradox” (McDermott, 2021).

The recent increase in usage and exploitation of freshwater resources is a consequence of rapid urbanization (Mall, et al., 2006). When combined with introducing alien species, it poses a danger to native biodiversity and ecosystem functioning. While many past research initiatives have improved our understanding of the negative impact of invasive species, there is still a fundamental void in the development of a robust and practical assay intended particularly for preliminary detection and monitoring (Kiruba *et al*.,2018). This vital gap is especially noticeable in India, where the distribution and spread of invasive fish species have not yet been thoroughly studied.

Biodiversity-rich India is still experiencing the continuous introduction of exotic fish species (Sandilyan, 2022) majorly due to two purposes, aquaculture and ornamental fish trade. But the potential impacts of these introductions on the environment, economy, and health are often not evaluated before importing (Bisen.,2023). Out of the 14 species listed as invasive by the National Biodiversity Authority (NBA) of India, 6 of them were introduced solely for the ornamental fish trade (Sandilyan, *et al.,2018*). Among those 6 listed species, four of them belong to the same genus, *Pterygoplichthys*, remarking it to be one of the most invasive genera of fish to be managed with immediate effect (Sandilyan *et al.,2018*).

The *Pterygoplichthys sp* (Armoured sailfin catfish.), native to South America, belongs to the largest catfish family *Loricariidae* (Order: Siluriform), and has a distinctive appearance with a long sail-like dorsal fin (10-12 fin rays), flat body structure, and bony armour of scutes covering whole body (Wu, et al., 2011). Their ventral suctorial mouth is specially designed to scrape off algal vegetation, hence, expected to be herbivores when introduced. However, recent studies have documented them to be omnivores feeding on eggs and dead carcasses (Bijukumar, *et al.,2015*). This fish was introduced in many countries, including India, because of their unique appearance and ability to manage tanks and aquariums from excess algae (Jumawan., 2016). They are now known for their remarkable size, high fecundity, and short-distance migratory behaviour outside the water, using their hard pectoral fins. They can tolerate oxygen-depleted water (Bijukumar, *et al.,2015*). Their razor-sharp fins and hard armour-like skin offer strong defence and make them less vulnerable to predators. Besides, the same structural design that gives them a defensive advantage also contributes to their reputation for causing damage to fishing nets (Chaichana & Jongphadungkiet., 2012). Their burrowing activity in dam walls and river banks for breeding also poses a serious risk of infrastructure collapse (Raj, *et al*., 2021). Hence, the detrimental effects of these catfishes go beyond the environment and have an economic impact on infrastructure and fisheries industries as well (Seshagiri, et al., 2021). This fish is not considered edible in most parts of India. However, concerted attempts are being made in a few places, like East Kolkata Wetland, to make them suitable for consumption. (Hussan *et al*.,2020). Fishermen often catch them as bycatch and dump them on the ground to decompose, which becomes a breeding ground for different flies and insects, creating a risk of various diseases for nearby villages and settlements (Clover., 2008).

The emergence of the invasive wild population of *Pterygoplichthys*, appears to result from unintentional release of aquarium specimens and escapees from overflown aquaculture farms during the rainy season. Favourable conditions in India, such as perennial sources of water, abundant food and promotions by aqua culturists as pets, allowed them to establish themselves as invasive. Species of the *Pterygoplichthys* genus are currently very difficult to distinguish due to high levels of interspecies hybridization found in the invading population. It has now become highly unreliable to distinguish these species based on their abdominal patterns (Bijukumar, *et al*.,2015). Hence, there is an urgent need for a molecular approach that can effectively target all species of this genus, as well as their presumed hybrids.

In recent years, the use of environmental DNA (eDNA) assays has gained popularity as a non-invasive method for detecting target species with accuracy and high sensitivity. Traditional methods like manual netting or visualization can be expensive, time-consuming, and harmful to endemic species (Dickie., *et al*., 2018). Instead, naturally released genetic material by an organism like faecal matter or sloughed cells are being used to detect genus/species-specific DNA barcodes, collected from water, soil sediment, or even air samples (Taberlet., *et al*., 2012). This approach not only aids to preserve the ecosystem but also makes it easier to detect invasive species, saving vital time and resources (Beng & Corlett., 2020) (Schmitter-Soto *et al*., 2016).

The Eastern Ghats, a discontinuous mountain range running on the east coastline boundary of India, is known for its unique biodiversity in terms of forest, water resources, and endemism of species (Subbaiyan *et al*., 2023). The freshwater fish diversity in this region is considered to be very high with many species of fish listed in the “world conservation union red list” (Sarkar & Uttam., 2006). Also, the list of identified species from this region is still growing, as new discoveries are made (Devi & Menon., 1995). For decades, the lakes and rivers of Eastern Ghats have been a source of sustenance for locals, with fishing being a vital component of their livelihood. This emphasizes the region’s importance as a hotspot for species richness and the required conservation efforts to maintain these valuable fish populations and their habitats. According to various research by Meena, et al., (2016), Raj, et al., (2020), Rao & Sunchu, (2017) and Bijukumar *et al*. (2015), many *Pterygoplichthys* species have been reported in various parts of India, notably in Kerala, Tamil Nadu, Andhra Pradesh, Uttar Pradesh, Bihar, and West Bengal. As these species are spreading at an alarming rate, there can be a possibility that they may have spread to many important and understudied sections of India, such as the Eastern Ghats. Researching the Eastern Ghats in relation to *Pterygoplichthys* invasion has become essential, as it allows us to comprehend the possibility of invasion risk, distribution, and ecological consequences of this fish species.

In this study, we aim to design a non-invasive, cost-effective, user-friendly and scalable tool for detecting and mapping invasive *Pterygoplichthys* species across freshwater ecosystems of the Eastern Ghats in India.

## 3. METHODS

### 3.1. DESIGNING OF PRIMERS

In response to the extensive hybridization found among the Indian population of *Pterygoplichthys* species (Bijukumar, *et al*.,2015), we opted to design a genus-specific primer, to identify existing *Pterygoplichthys* and their hybrids in freshwater ecosystems.

For this, we focused on the ‘Cytochrome b’ region of the mitochondria to develop primers. We obtained sequences from NCBI GenBank for four species from different genera: *P. pardalis* (NC_060468.1:14362-15499), *P. disjunctivus* (NC_015747.1:14362-15499), *Hypostomus plecostomus* (NC_025584.1:14362-15499), and *Wallago attu* (NC_063564.1:14363-15500). Here, *P. pardalis* and *P. disjunctivus* belong to our target genus, while *Hypostomus* was a phylogenetically closely related genus from South America sharing the same native region as *Pterygoplichthys*, whereas *Wallago* is a phylogenetically distantly related genus of catfish commonly found in Indian water bodies. Upon obtaining sequences from the database, they were aligned using the MUSCLE (Multiple Sequence Comparison by Log-Expectation)-based method in Clustal Omega (https://www.ebi.ac.uk/Tools/msa/clustalo/). The alignment was visually examined to identify a region that exhibited high variability among the Genera while demonstrating low variation within the species of the target genus. Using the same alignment as input, we developed a genus-specific primer using IDT’s PrimerQuest Tool (https://www.idtdna.com/) (see supplementary file for primer parameters). The resultant primer pairs were meticulously evaluated in IDT’s OligoAnalyzer™ Tool (https://www.idtdna.com/calc/analyzer), to assess their propensity for primer dimers, secondary structure formation, melting temperature, and amplicon size. With the help of NCBI’s Primer-Blast tool, we performed in-silico screening to ensure that the designed primers were genus-specific.

### 3.2. IN VITRO ANALYSIS OF PRIMERS

To ensure the specificity of primers, it is essential to complement the in-silico analysis with in-vitro validations. DNA databases are still growing and may not contain sequences of many understudied or recently described species. Hence to test the specificity of our primer pair in-vitro against the genus of *Pterygoplichthys*, Genomic DNA was obtained from tissue samples of 3 species of our target genus “*Pterygoplichthys”* Particularly *P. disjunctivus* and *P. pardalis* were collected from wild whereas *P. joselimaianus* was procured from an ornamental fish farm. . A total of 33 non-target species of fish, including 19 catfish (Siluriform) and 4 non-catfish species co-existing with invasive residents of *Pterygoplichthys* in Indian aquatic habitats (sympatric) and 7 other fish species that inhabit the same native environment as *Pterygoplichthys* in South America (Allopatric) were chosen for this study. Prior to that, an additional step for species confirmation using a COI marker was also performed according to Ivanova, et al.’s 2007 paper. The comprehensive list of species used for assessing primer specificity is given in table-1 below.

**Table 1:** List of species considered for the Invitro primer specificity testing, which includes phylogenetically distantly related species sharing the same ecosystem in India as well as phylogenetically closely related species sharing native range in South America.

| **Siluriform Species from India** |  |  |
| --- | --- | --- |
| *Pangasianodon hypophthalmus*<br>(Pangasiidae) | *Arius arius*<br>(Ariidae) | *Lepidocephalichthys guntea*<br>(Cobitidae) |
| *Mystus vittatus*<br>(Bagridae) | *Pseudolaguvia shawi*<br>(Erethistidae) | *Botia dario*<br>(Botiidae) |
| *Pachypterus atherinoides*<br>(Horabagridae/ Schilbidae) | *Chaca chaca*<br>(Chacidae) | *Sisor rabdophorus*<br>(Sisoridae) |
| *Heteropneustes fossilis*<br>(Heteropneustidae) | *Hara jerdoni*<br>(Erethistidae) | *Ompok bimaculatus*<br>(Siluridae) |
| *Erethistes pusillusi*<br>(Erethistidae) | *Olyra longicaudata*<br>(Bagridae), | *Bagarius bagarius*<br>(Sisoridae) |
| *Amblyceps mangois*<br>(Amblycipitidae) | *Sperata aor*<br>(Bagridae) | *Clarias dussumieri*<br>(Clariidae) |
| *Wallago attu*<br>(Siluridae) |  |  |
| **Non- Siluriform Species from India** |  |  |
| *Labeo rohita*<br>(Cyprinidae) | *Tenualosa Ilisha*<br>(Clupeidae) | *Labeo catla*<br>(Cyprinidae) |
| *Oreochromis mossambicus*<br>(Cichlid) |  |  |
| **Species Sharing Common Native Range in South America** |  |  |
| *Corydoras paleatus*<br>(Callichthyidae) | *Platydoras armatulus*<br>(Doradidae) | *Synodontis eupterus*<br>(Mochokidae) |
| *Ancistrus temminckii*<br>(Loricariidae) | *Trachelyopterus galeatus*<br>(Auchenipteridae) | *Crossocheilus atrilimes*<br>(Cyprinidae) |
| *Hypancistrus zebra*<br>(Loricariidae) |  |  |

During primer validation, a PCR reaction was set up with of a total volume of 15μl/reaction., which includes 4.6μl Nuclease free water, 7.5μl of EmeraldAmp GT PCR 2X Master Mix, 0.2μl of forward and 0.2μl of reverse primer (10 μM), 1.5μl of 0.1% BSA, and 1μl of template DNA. The cycler was programmed with an initial activation step at 94 °C for 5 minutes, followed by 40 cycles of PCR with denaturation at 94 °C for 30 seconds, annealing at 56 °C for 20 seconds, extension at 72 °C for 30 seconds, and final extension step at 72 °C for 5 minutes. The resulting PCR products were visualized using 2% agarose gel and were validated by sequencing.

### 3.3. qPCR STANDARD PREPARATION

To accurately estimate the copy number of the target DNA in field samples, the use of a standard curve as a reference is essential. For preparing the standard a bulk PCR was performed in 100μl (5 reactions of 20μl each) reaction using tissue-extracted DNA from *Pterygoplichthys* sp. as the template to amplify a 163 bp fragment of the Cytochrome B gene using the designed primers, confirming the amplification using 2% agarose gel. Upon confirmation, the amplified products were purified using the Nucleospin® Gel and PCR Clean-up kit, according to the manufacturer’s instructions and quantified using a nanodrop spectrophotometer. The DNA concentration in ng/μl and the amplicon size were used as input in the online calculator (https://cels.uri.edu/gsc/cndna.html) to calculate the copy number. A 10-fold serial dilution was performed using the purified PCR product to prepare standards ranging from 10^5^ to1 copies/μl to generate the standard curve. For the Limit of detection (LOD) and limit of quantification (LOQ), a qPCR was performed using the same purified PCR product but with, 4-fold dilutions ranging from 1024 copies/reaction to 1 copy/reaction with 12 technical replicates and calculated using LOD/LOQ Calculator R Script (Klymus., *et al*.,2020a). The results were calculated in copy numbers.

For qPCR, the total reaction volume was set to be 10μl which includes 3.6μl DNase free water, 0.2μl of 10 μM forward and reverse primers, and 5μl of TB Green® Premix Ex Taq™ II—Tli RNaseH Plus and 1μl of the required standard. The qPCR was performed with Roche Light cycler 480 II using the conditions as denaturation at 95 °C for 5 seconds followed by 40 cycles of PCR at 62°C for 5 seconds, then melting step starting from 60 °C for 60 sec with a progressive rise in temperature till it reaches 95 °C with a slow ramp rate of 1.1 and finally cooling at 50 °C for 30 seconds.

### 3.4. FIELD SAMPLING

We selected 20 Waterbodies including lakes, reservoirs, and dams in and around the Eastern Ghats spanning the states of Tamil Nadu, Telangana, Andhra Pradesh, and Odisha (Figure 1) to investigate the presence of *Pterygoplichthys sp*. via our assay. The selection of water bodies was based on a few considerations, notably the existence of perennial water, proximity towards significant wildlife, connectivity via roadways, and places where aquaculture is regularly practised. For sampling, we used Smith root environmental DNA sampler for filtering water using Mixed Cellulose Ester (MCE) as a filter membrane of pore size 0.45 microns (Thomas., *et al*., 2018). Among the 20 locations that were selected, each water body was sampled at multiple locations in triplicates. Depending on the turbidity of the water, the volume of water filtered from each water body ranged from 3 to 6 litres. The sampled filter papers were stored in the same self-preserving filter assemblies.

**Figure 1:**
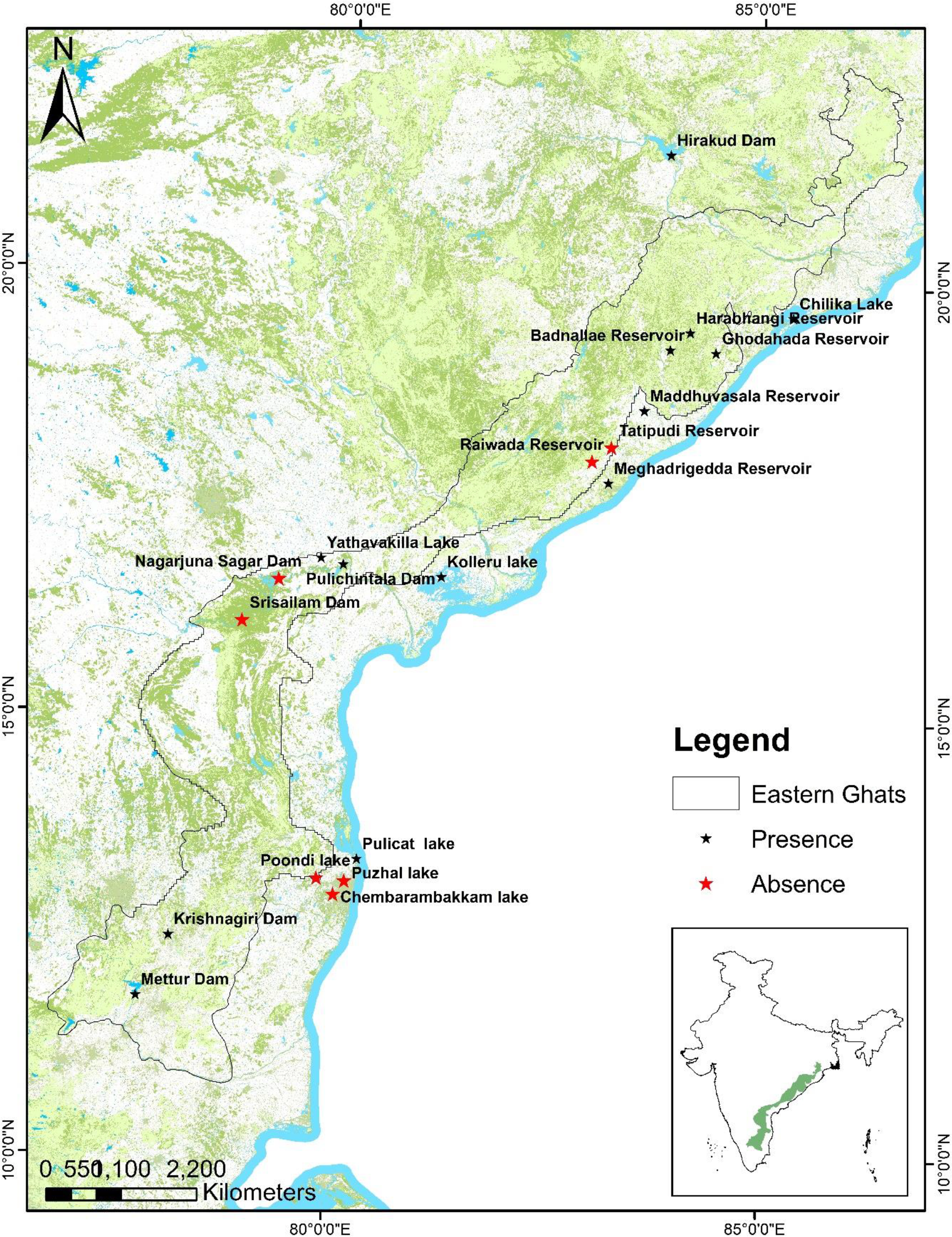
The Eastern Ghats map showing sample collection locations and positive and negative for *Pterygoplichthys* determined by the developed eDNA assay

Along with eDNA sampling, with the help of local fishermen we also conducted a short survey for the confirmation of species in the same waterbodies (see supplementary file). After sampling the filter membranes were brought back to the lab and immediately isolated batch-wise along with extraction negative control, using Qiagen Blood and Tissue Kit. For the environmental positive control (EPC) we sampled eDNA from a pond neighbouring the laboratory campus, where the presence of *Pterygoplichthys sp*. was visually confirmed, while eDNA samples obtained from our laboratory fish tank, which was free of *Pterygoplichthys sp*. was considered as environmental negative control. The extracted DNA concentration (see supplementary file) was determined using Thermo Fisher Scientific’s Qubit Fluorometers and preserved at −20 °C till further analysis.

### 3.5. DETECTING THE PRESENCE OF *PTERYGOPLICHTHYS SP*. IN eDNA SAMPLES

DNA concentrations of all samples were adjusted to 10ng/μl in order to avoid sample inhibition in the assay. A small fraction of the sample was already less than the concentration of 10ng/μl and hence was not adjusted. Each sample was loaded in the qPCR plate as technical triplicates, with the same reaction volume and conditions as described during the optimization of qPCR. Those reactions which didn’t show any amplification or had primer dimers were checked for target in secondary peaks of melting curve analysis and tested again. Copy numbers were calculated using the generated standard curve for reactions that showed amplification. In order to confirm the assay results, amplified qPCR products were purified using NucleoSpin® Gel and PCR Clean-up (Macherey–Nagel, Germany) kit and sequenced via ABI 3730XL DNA sequencer platform using the BigDye Terminator (version 3.1) Cycle Sequencing Kit and POP-7 Polymer separation matrix (Applied Biosystems, Inc.) using both forward and reverse primer. To obtain consensus sequences, we used CodonCode Aligner software (version 8.0) and the resultant contigs were analysed against the NCBI’s nucleotide BLAST to verify species. All the steps of this assay were done carefully in a laboratory dedicated to qPCR work, to avoid cross-contamination.

## 4. RESULTS

### 4.1. SCREENING OF PRIMER PAIR

Adhering to all the parameters of primer designing and screening them both in-silico, we designed a primer pair of 163bp amplicon size, targeting the mitochondrial Cytochrome-B region of Pterygoplichthys species. The list of designed primer pair along with its sequence is given in Table-2.

**Table 2:**
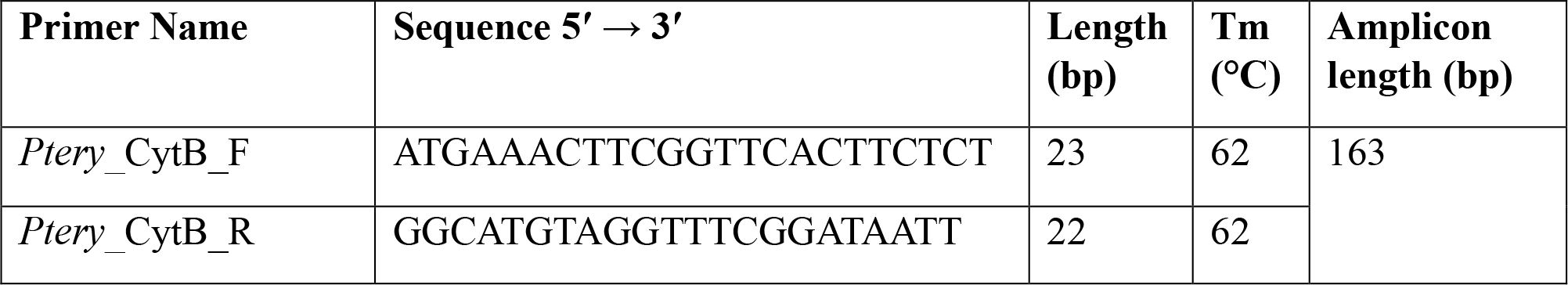
Designed primer pair and their sequence used for the detection of Pterygoplichthys sp.

### 4.2. IN VITRO PRIMER SPECIFICITY ASSAY

Primer specificity testing revealed prominent bands exclusively with *Pterygoplichthys* species at 163bp. No amplification was observed for any of the non-target species included in the study (Figure 2). Additionally, the identity of all the species included in this assay was confirmed using the universal COI marker (see supplementary file). In both cases, no bands were observed in the negative control.

**Figure 2:**
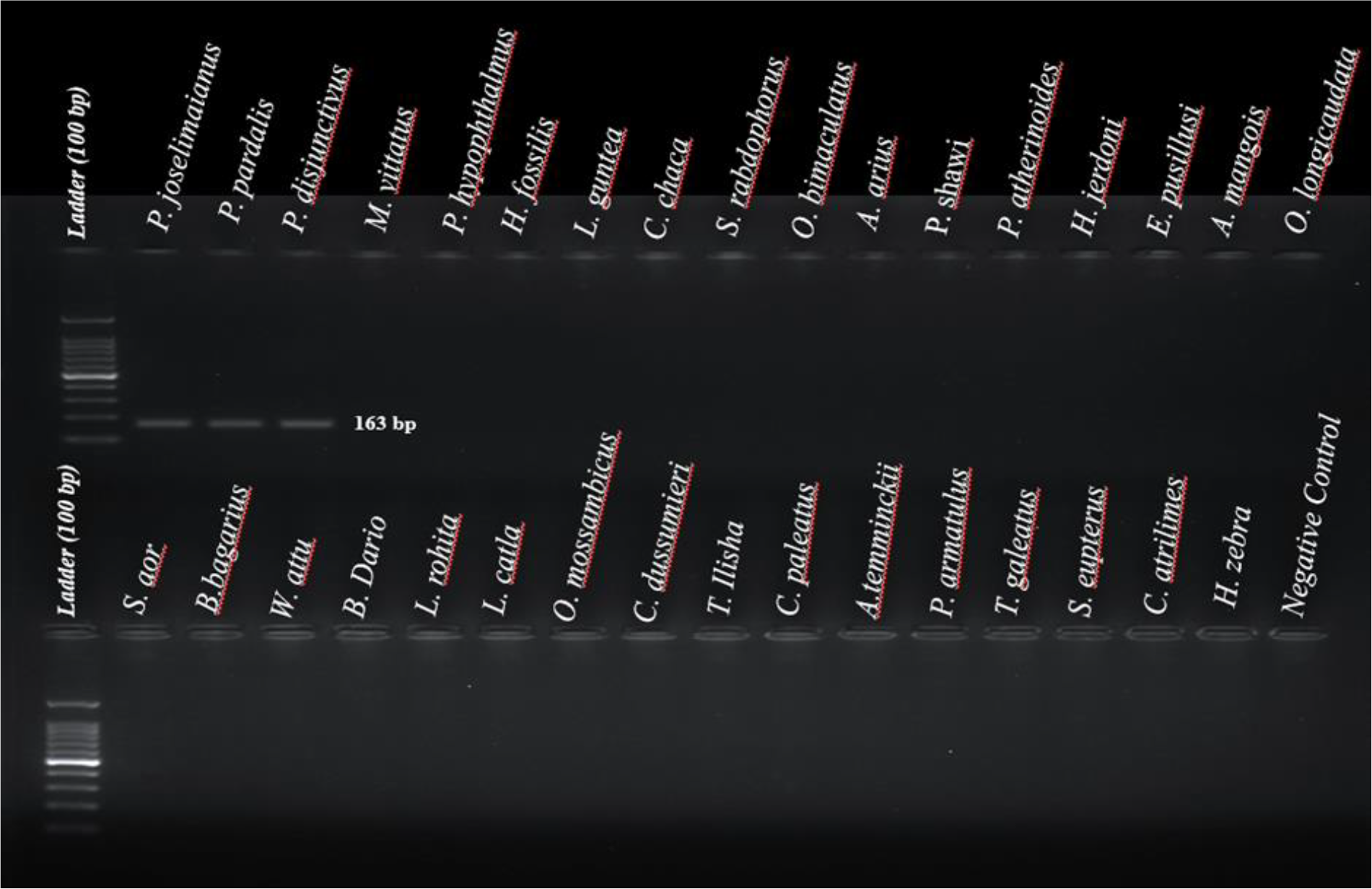
Agarose Gel picture representing the in-vitro PCR validation of designed primers against the three species of Pterygoplichthys and 30 additional non-target species.

### 4.3. qPCR STANDARDISATION

The standard curve prepared with three technical replicates for each dilution showed an efficiency of 97.99% with an R^2^ value of 0.9978. The predicted Ct of a reaction with just a single copy of the target (y-intercept) was found to be 34.546 (Figure 2). For the sensitivity of the assay, the limit of detection (LOD) was calculated to be 4 copies and the limit of quantification (LOQ) was found to be 12 copies.

### 4.4. *Pterygoplichthys* DETECTION IN FIELD eDNA SAMPLES

This assay, intended to detect *Pterygoplichthys*, has shown significant findings in a variety of locations. In Tamil Nadu, three of the six sampled locations showed positive results. The same result was reported from two out of four locations in Telangana, however in Andhra Pradesh, the assay revealed the presence in three out of the five locations analysed. Interestingly, all the eDNA samples collected from Odisha were tested to be positive during the assay (Figure 1 and 4). These outcomes suggest a widespread distribution of *Pterygoplichthys* in the studied areas across the eastern ghats. It was noticed that the PCR triplicates within each field replicate produced consistent findings, along with the Environmental Positive Sample (EPC) regularly showing positive amplification, demonstrating the assay’s dependability. There was no amplification in the environmental negative samples (ENC) or the extraction blanks, validating the absence of false positives. Using the standard curve and the qPCR-generated CP (Cycle threshold) value, the copy number for each positive sample was calculated. It was observed that all positive samples exhibited copy numbers surpassing the calculated limits of detection (LOD) and quantification (LOQ) established for the assay. It was found that the sequence had a significant similarity with *Pterygoplichthys pardalis* and *Pterygoplichthys disjunctivus*. The percentage identity of the matches varied from 95% to 100%, with query coverage ranging from 98% to 100%.

**Figure 3:**
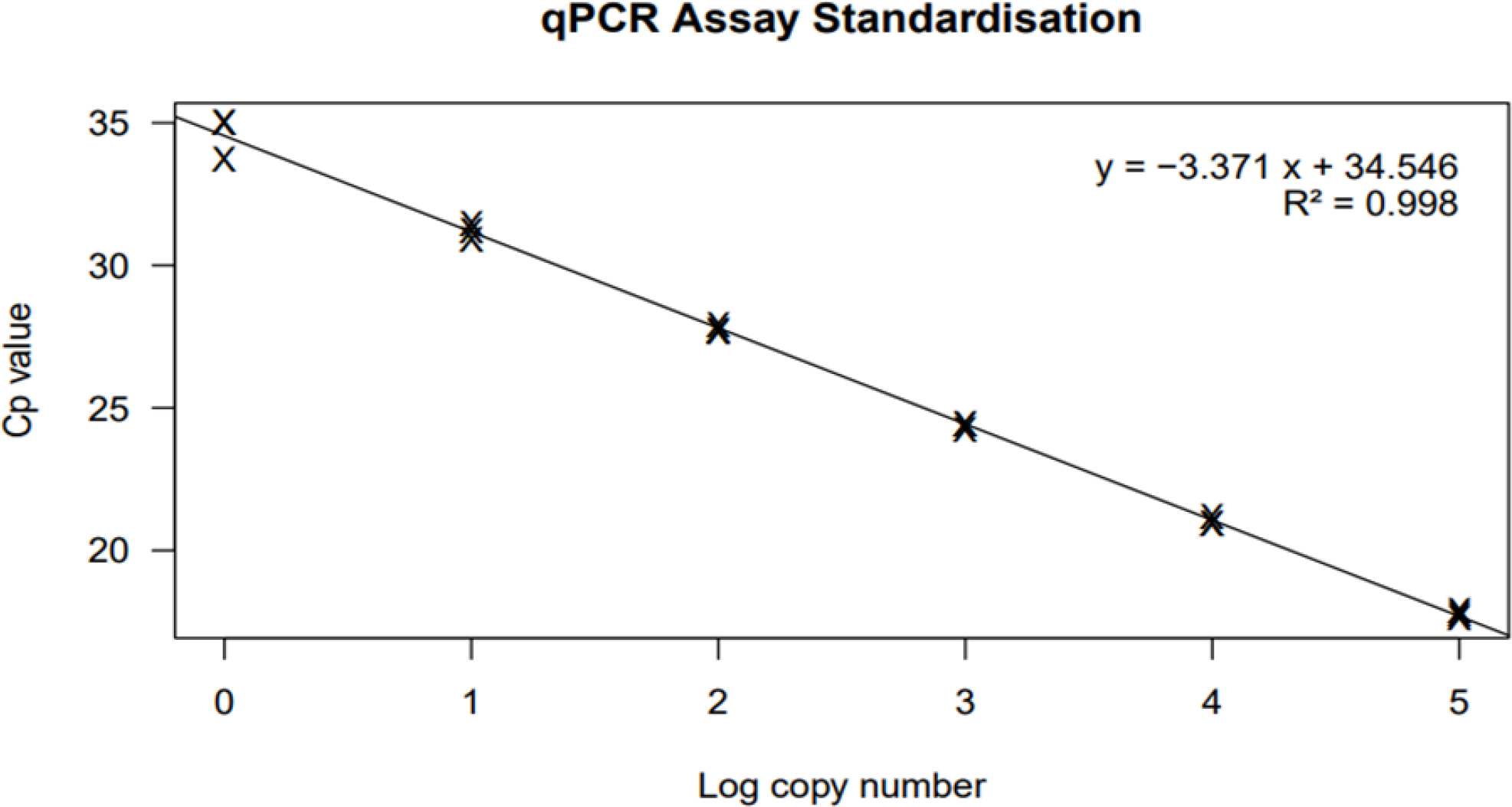
The generated standard curve derived from the amplification of target DNA with a 10-fold dilution of concentration ranging from 1 to 105 copies.

**Figure 4:**
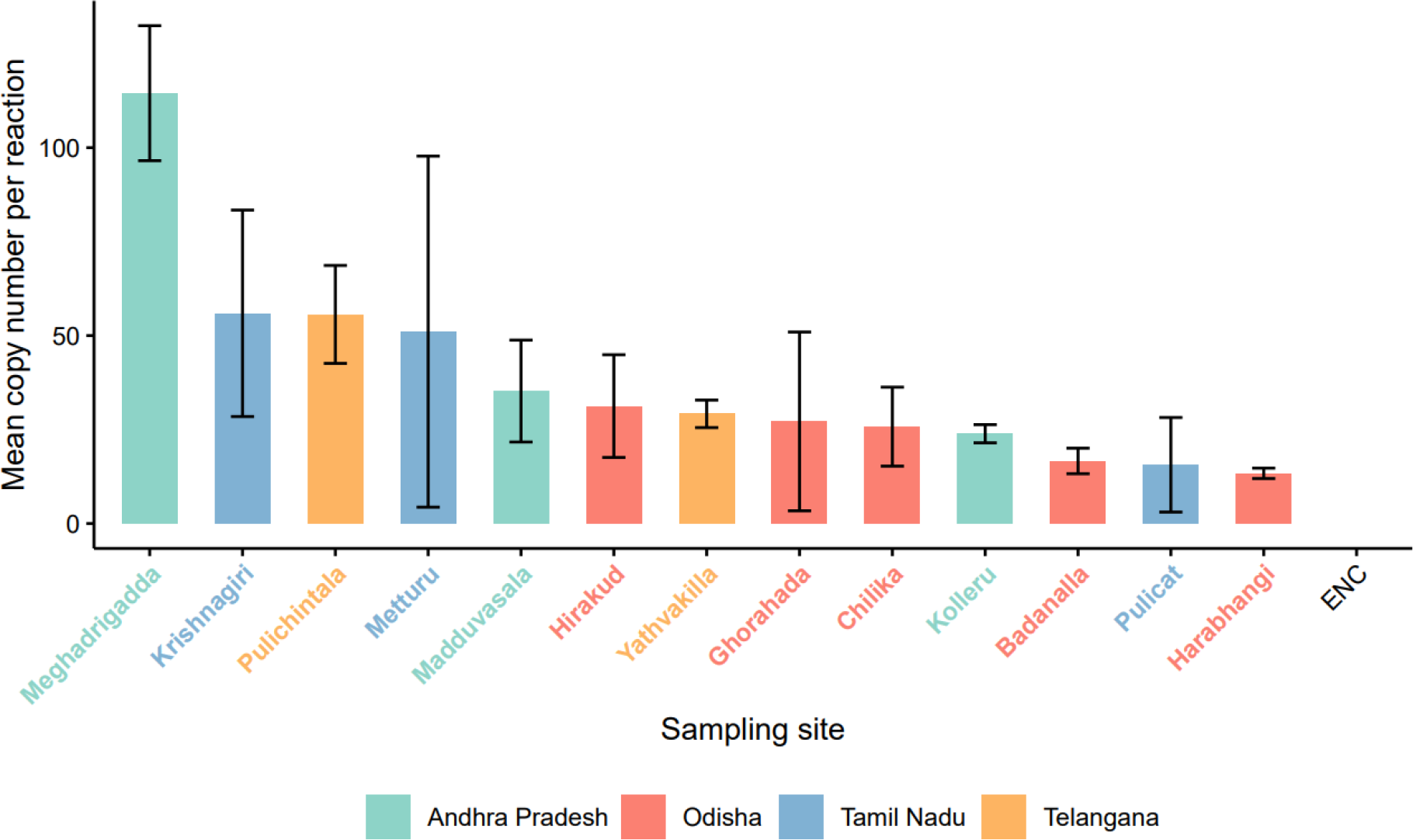
Bar graph representing mean copy number/reaction for different locations that showed positive for *Pterygoplicthys* via this eDNA assay

## 5. DISCUSSION

The presence of invasive *Pterygoplichthys* in the Eastern Ghats is a major concern owing to the region’s diverse aquatic ecology which supports many native fish species. In particular, Andhra Pradesh and Telangana have been found to support 152 freshwater fishes (Krishna & Srinivasulu., 2021), Odisha comprises 186 species (Mogalekar & Johnson.,2018b), and Tamil Nadu has an impressive count of around 226 species (Mogalekar & Johnson.,2018a). The invasion of *Pterygoplichthys* is considered a serious risk, due to their habit of mass consumption of algae and detritus, disrupting the natural nutrient cycle, and contributing to ecological imbalances along with water quality issues (Clover., 2008). Hybridization has further complicated the identification of these species. With each passing day revealing new records of its presence in various water bodies, it has become highly essential to detect its presence as early as possible for proper management.

Existing studies in India by Dharan et al. (2015), as well as in Southeastern Asia by Page and Robins (2006), have demonstrated the significance of invasive *Pterygoplichthys* monitoring. However, a majority of research involving monitoring is still based on traditional methods, which have drawbacks. Although methods like electrofishing, netting, and visual surveys are widely used, they can provide difficulties in the proper identification and quantification of species, especially when dealing with hybrids or larval stages of fishes. Complete dependency on such methodologies can lead to false negatives and wrong interpretations. Conventional techniques are often restricted to smaller geographical coverage because of their high time and labour-intensive nature. As a result, there is an increasing demand for incorporating sophisticated molecular-based techniques.

To address this need, we could develop exceptionally reliable and specific qPCR-based detection assays for invasive populations of *Pterygoplichthys* species found in India. Since the primary aim is to target both pure and hybrid species of *Pterygoplichthys* without distinguishing them, the use of the mitochondrial Cytochrome B gene as a marker solves this problem as mitochondria are always inherited maternally. Another reason is the relatively high copy number within a cell compared to any other nuclear gene, hence increasing the chances of detection. Our assay design, testing, and validation processes line up with Goldberg et al. (2016) recommended parameters for eDNA assay development. The specificity of the marker and the sensitivity of the assay were considered to be the two important components of this assay (Klymus., *et al*., 2020b). During specificity checking of the marker in-silico, the primer was also found to target *Baryancistrus xanthellus*, from the same Loricariid family of catfish. This may be due to the fact that Loricariidae being the biggest family of catfish, contains many subspecies which are genetically very close. Since *Baryancistrus xanthellus* is completely exotic in India and its presence in Indian waterbodies would definitely be considered invasive so this was not considered a major issue during the designing of primers. Even for invitro specificity testing of primers, this marker could clearly identify all three species of *Pterygoplichthys* considered for this study, namely *P. pardalis, P. disjunctivus*, and *P. joselimaianus*. Hence, we strongly advise validating the marker on other *Pterygoplichthys* species not included in this study to ensure the assay’s consistency. It would also be beneficial to include other genetically comparable species within their geographic area of interest during reproducing this assay in other countries apart from India. In case of sensitivity, this assay has proven to be effective in recognizing barely detectable levels of *Pterygoplichthys* eDNA, with detection limits as low as 4 target copies per reaction. Since the primer designed for this study was found to be specific for *Pterygoplichthys* species, we refrained from the use of any probe. This made the assay simpler and cheaper to reproduce.

The assay proved capable of detecting *Pterygoplichthys* even in locations where no visual findings had been documented. This emphasizes the assay’s potential to detect the target species even at an early stage of invasion, outperforming traditional detection approaches.

However, Invasive species management needs periodic surveillance. Although our designed assay was effective in detecting *Pterygoplichthys*, findings from just one-time sampling may not be adequate. Hence, seasonal sampling of the waterbodies would considerably improve the effectiveness of the monitoring process. Invasion is a global concern, with increasing public awareness of invasive species and employing assays like this for management will substantially contribute to efficiently addressing the invasion issue. Our assay can definitely aid with the monitoring stages of the management approach. This can provide a validation for prompt action, by conservation officials and policymakers, to make sensible judgments regarding the management of *Pterygoplichthys* in the Eastern Ghats. Not only that, the findings might promote additional research in specific fields like invasion pathways and patterns, impact assessments, the study of hybridization, and food web research, all of which will aid in our better understanding of this species. However, avoiding fresh invasions is highly essential for minimizing the damage to this rich and vulnerable environment.

The escalating proliferation of *Pterygoplichthys* in the Eastern Ghats presents a challenge to native fish diversity because of their competition for food and shelter. Considering the checklist on the fish diversity of the eastern ghats prepared by Lakra & Sarkar in 2006 revealed the presence of several species listed on the World Conservation Union Red List including the Deccan mahseer (*Tor khudree*). However, the current status of endemism in this region still remains unclear due to the lack of current studies. Identifying the presence of *Pterygoplichthys* via eDNA assay in internationally important sites (Ramsar sites) like Chilika lake and Hirakud Dam in Odisha and Kolleru lake in Andhra Pradesh has accelerated the apprehension by many folds on the survival of endemic and endangered species present there. In such a situation early detection of invasive *Pterygoplichthys* holds immense importance, as an increase in dominance by *Pterygoplichthys* can displace or even eliminate endemic species that might not yet be fully understood or documented. This assay not only offers a reliable method for monitoring invasive species but also lays the foundation for strategic management initiatives, by offering relative quantification between different study areas. This would be a crucial step toward prioritizing waterbodies that require immediate attention and management against invasion, without disturbing the areas that are still not invaded by *Pterygoplichthys*.

In a nutshell, the prior detection of *Pterygoplichthys* is crucial for preserving ecosystems and supporting ecological equilibrium. The study achieved its goal by offering a dependable technique for early detection of *Pterygoplichthys* distribution in India’s Eastern Ghats, which greatly adds to continuing invasive species management efforts. Our assay is a very useful initial step toward mapping the present invasion and managing *Pterygoplichthys*, with the aim of conserving our native species wherever feasible.

## Supplementary Information

### 1. Parameters for primer designing in IDT DNA-Primer Quest

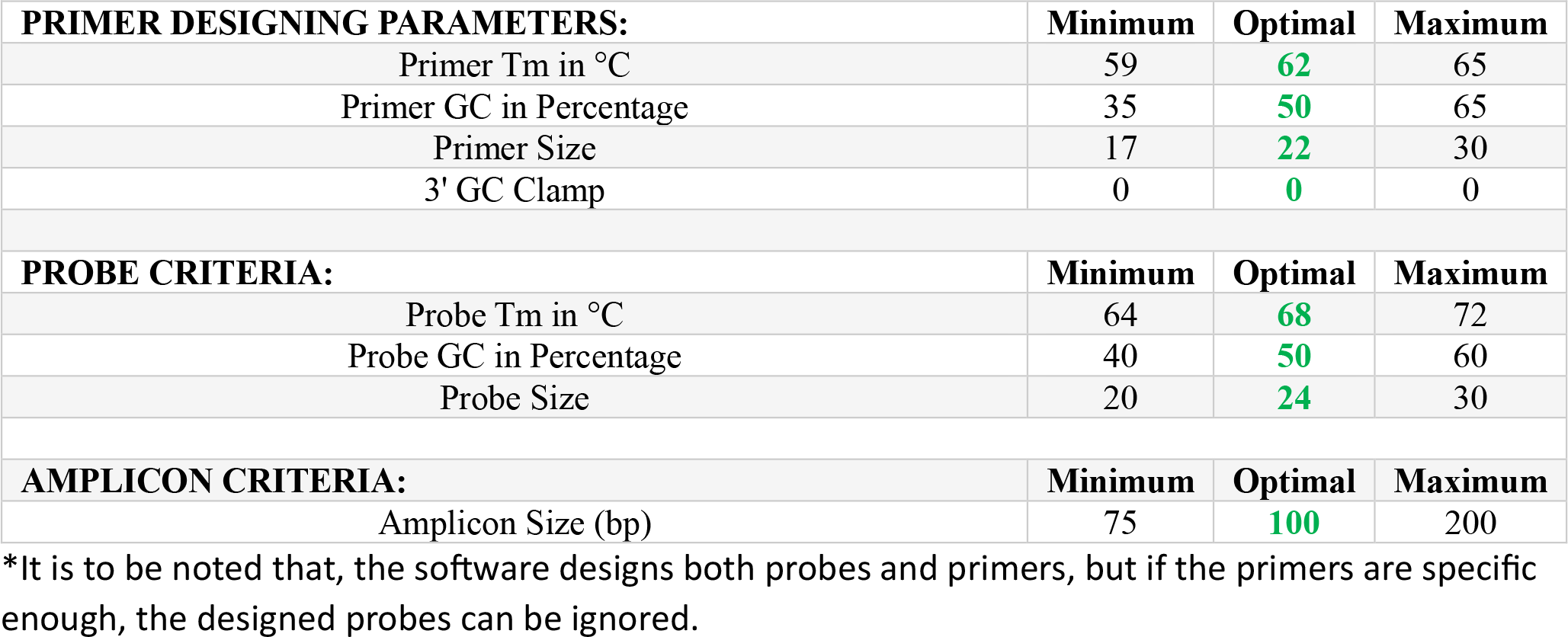

### 2. Species Confirmation for Invitro primer specificity testing

**Figure:**
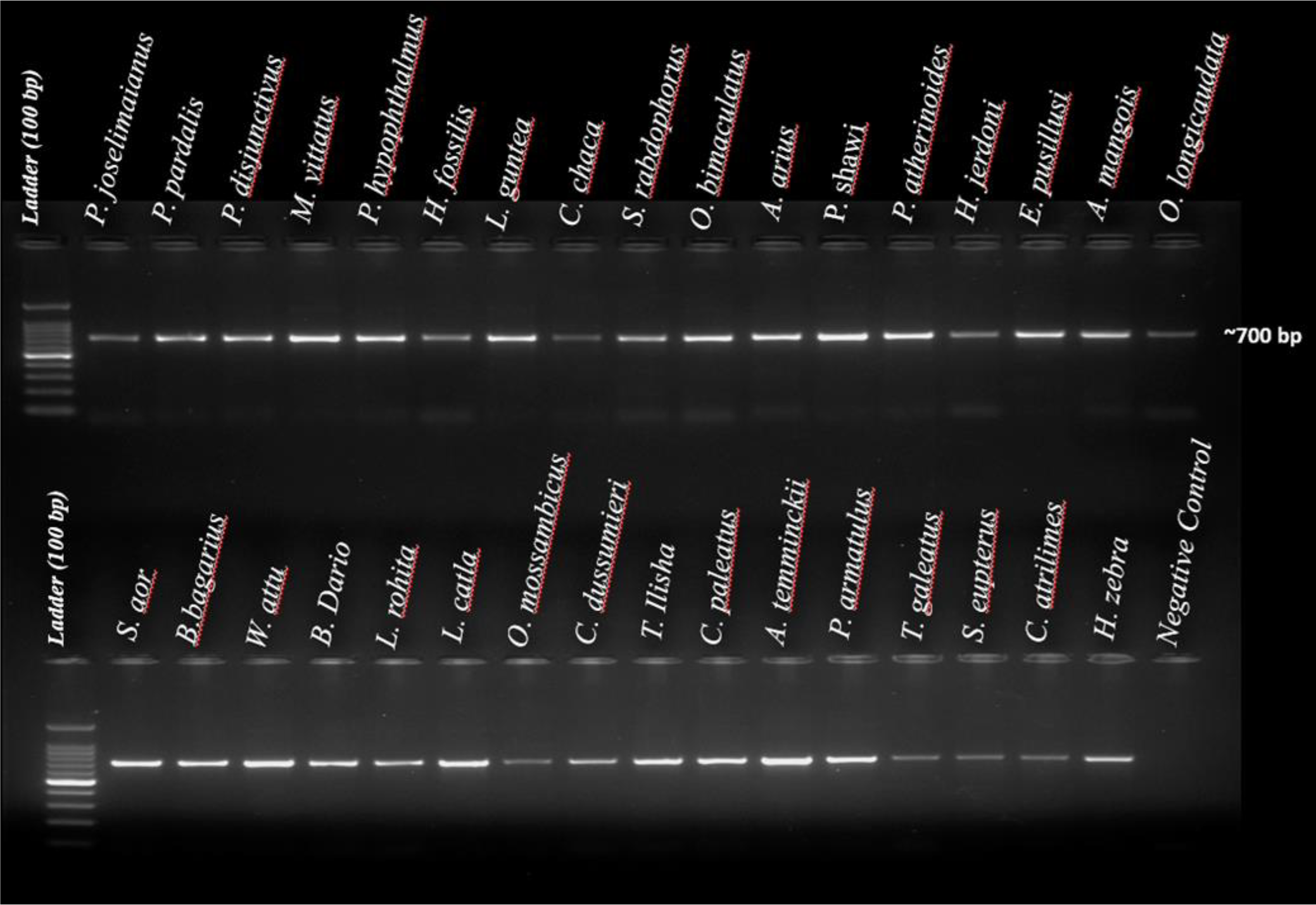
Gel picture showing, species confirmation using COI marker, for all species considered for Invitro primer specificity of the designed primers (Ivanova, N. V., et al., 2007).

### 3. Additional Data regarding sampled locations across the Eastern Ghats of India

**Table:**
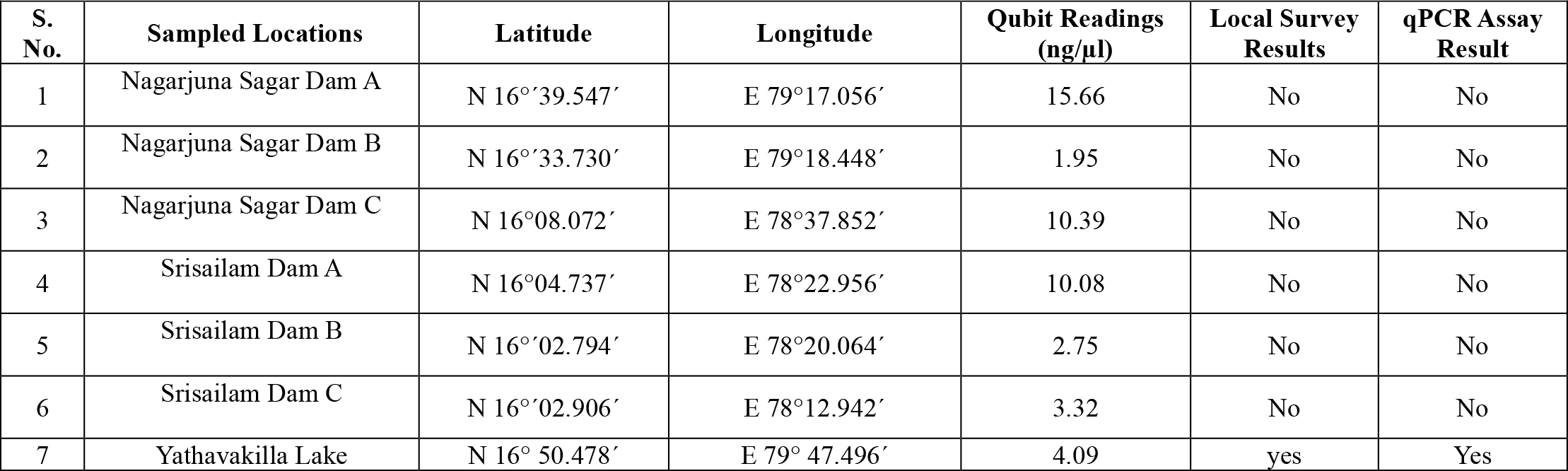
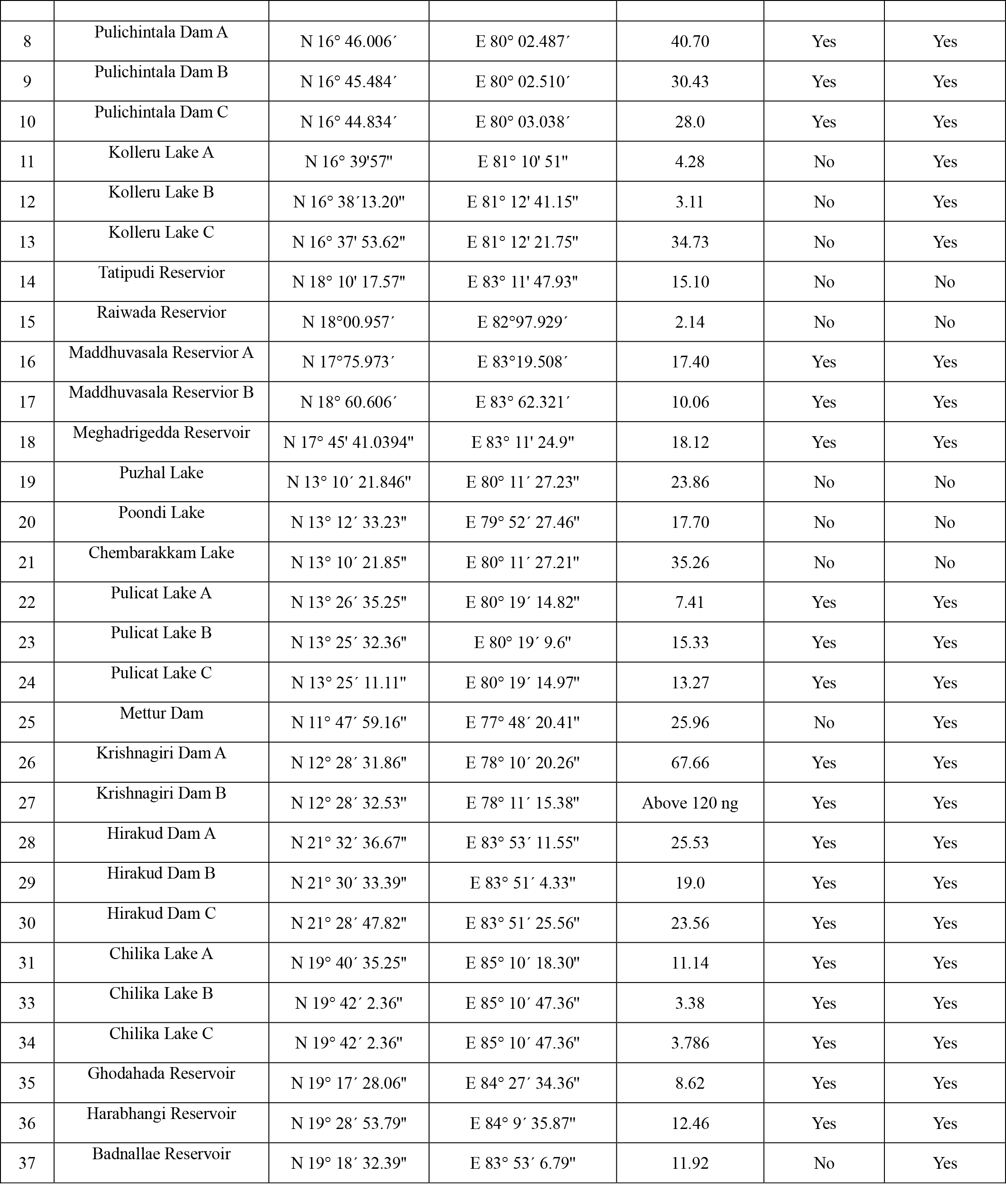
Additional Information Including Location coordinates, Local Fisher man survey results and the qPCR assay results of the eDNA sampled locations across the eastern ghats of India.

